# Evaluating the Significance of Embedding-Based Protein Sequence Alignment with Clustering and Double Dynamic Programming for Remote Homology

**DOI:** 10.1101/2025.07.28.666913

**Authors:** Robert Spicer, Nilanjana Raychawdhary, Cheryl Seals, Sutanu Bhattacharya

## Abstract

Accurate detection of protein sequence homology is essential for understanding evolutionary relationships and predicting protein functions, particularly for detecting remote homology in the “twilight zone” (20-35% sequence similarity), where traditional sequence alignment methods often fail. Recent studies show that embeddings from protein language models (pLM) can improve remote homology detection over traditional methods. Alignment-based approaches, such as those combining pLMs with dynamic programming alignment, further improve performance but often suffer from noise in the resulting similarity matrices. To address this, we evaluate a newly developed embedding-based sequence alignment approach that refines residue-level embedding similarity using K-means clustering and double dynamic programming (DDP). We show that the incorporation of clustering and DDP contributes substantially to the improved performance in detecting remote homology. Experimental results demonstrate that our approach outperforms both traditional and state-of-the-art approaches based on pLMs on several benchmarks. Our study illustrates embedding-based alignment refined with clustering and DDP offers a powerful approach for identifying remote homology, with potential to evolve further as pLMs continue to advance.

## Introduction

Identifying protein sequence homology through sequence similarity remains a standard approach for detecting evolutionary conserved functions across proteins^1,2^. For decades, protein sequence homology supports numerous applications, including the prediction of protein functions^3–7^, protein structures and protein interactions^8–17^, protein design^18^, and evolutionary relationships^1^. Traditional sequence homology-based methods are typically fast and accurate when proteins share high sequence similarity. However, while protein sequence similarity is usually below 20-35%, often referred to as the twilight zone^19^, their accuracy declines rapidly. It is well-established that protein structure is more conserved than sequence across evolutionary time^20^, making remote homology detection, the task of detecting structurally similar proteins with low sequence similarity, a major challenge for existing sequence homology-based approaches.

Moreover, structure-based alignments tools such as TM-align^20^, Dali^21^, FAST^22^, and Mammoth^23^ can accurately detect such remote homologs by superimposing protein three-dimensional (3D) structures. Nonetheless, they need experimentally determined or predicted structures that remain unavailable for most proteins. Despite the recent progress made by protein structure prediction methods^9,11,12,24,25^, including AlphaFold2^10^, that transforms the field with rapid structure predictions with high accuracy, it still faces critical limitations, particularly in keeping pace with the exponential growth of number of available protein sequences^26^. For example, metagenomic alone has billions of unique protein sequences^27,28^, of which only a small fraction has known structures^29^, thereby highlighting the importance of efficient sequence-based approaches that capture structural similarities without requiring explicit structure prediction.

Recent advances in sequence-based approaches using pre-trained protein language models (pLM) show promise. These transformer-based models, inspired by advances in natural language processing, are trained on millions of protein sequences using self-supervised learning. These models treat protein sequences as sentences. Through this training, pLMs begin to understand the “language of life” by capturing important biological information^30^. When a protein sequence is input, pLM produces high-dimensional vector representations, known as embeddings, for each residue or for the entire sequence. In recent years, these embeddings become an important feature for sequence alignments, particularly for remotely homologous proteins in the twilight zone. To do this, several methods represent protein sequences by averaging residue-level embeddings into fixed-length vectors in high-dimensional space^31–33^. Evo-velocity^32^ uses Euclidean distance between these averaged embeddings to construct evolutionary graphs via K-nearest neighbors, while ProtTucker^31^ employs contrastive learning to improve clustering of similar CATH domains^34^. TM-Vec^29^ also uses averaged embeddings to directly predict TM-scores^35^ for computing structural similarity. Although these approaches are efficient, they often overlook fine-grained residue-level alignment information^36^. To overcome this, alignment-based strategies^37–39^ are introduced, which compute residue-residue similarity using embedding-derived alignments or train neural networks to produce scoring and gap parameters. Most recently, EBA^36^ proposes an unsupervised alignment-based approach that combines residue-level embedding similarities with dynamic programming to detect structural relationships in the twilight zone, outperforming existing pLM based approaches. However, alignment-based methods suffer from noise in the resulting similarity matrix^36^.

So, can these resulting embedding similarity matrices be refined in an unsupervised manner to effectively detect remote homology? To address this challenge, we develop a new unsupervised protein sequence alignment approach that refines residue-level embedding similarity by incorporating K-means clustering and a double dynamic programming strategy. We evaluate the effectiveness of our approach through a threefold strategy. First, we perform structural alignment benchmarking on the PISCES dataset^40^ (≤ 30% sequence similarity) by calculating Spearman correlations between predicted alignment scores and TM-align^20^–derived similarity scores (TM-scores) to evaluate structural similarity across remote homologs. Second, we conduct an ablation study to assess the contribution of each component in our pipeline— specifically, the clustering-and double dynamic programming-based refinement introduced in this work—by systematically removing these components and measuring their individual effects on alignment performance. Third, we evaluate functional generalization using CATH^34^ annotation transfer task across all classification hierarchy (Class, Architecture, Topology, and Homology). Notably, our approach requires no training or parameter optimization, is compatible with multiple pretrained embeddings, making it inherently interpretable and robust.

## Methods

### Protein Embeddings from Language Models

As shown in **Figure 1**, the initial step of our approach converts protein sequences into residue-level embeddings using pretrained protein language models (pLMs). These models generate high-dimensional vector representations for each residue, capturing both sequence context and physicochemical properties. In this study, we use residue-level embeddings generated by three widely used pLMs: ProtT5 (ProtT5-XL-UniRef50)^30^, ProstT5^41^, and ESM-1b (esm1b_t33_650M_UR50S)^42^. While ProtT5 and ESM-1b are transformer^43^-based models trained on the UniRef50^44^ database using masked language modeling, ProstT5 builds upon ProtT5 by incorporating sequential and structural information through Foldseek^45^’s 3Di-token encoding. These models output fixed-length vectors for each residue, with dimensionalities of 1024 for ProtT5 and ProstT5, and 1280 for ESM-1b.

**Figure 1.**
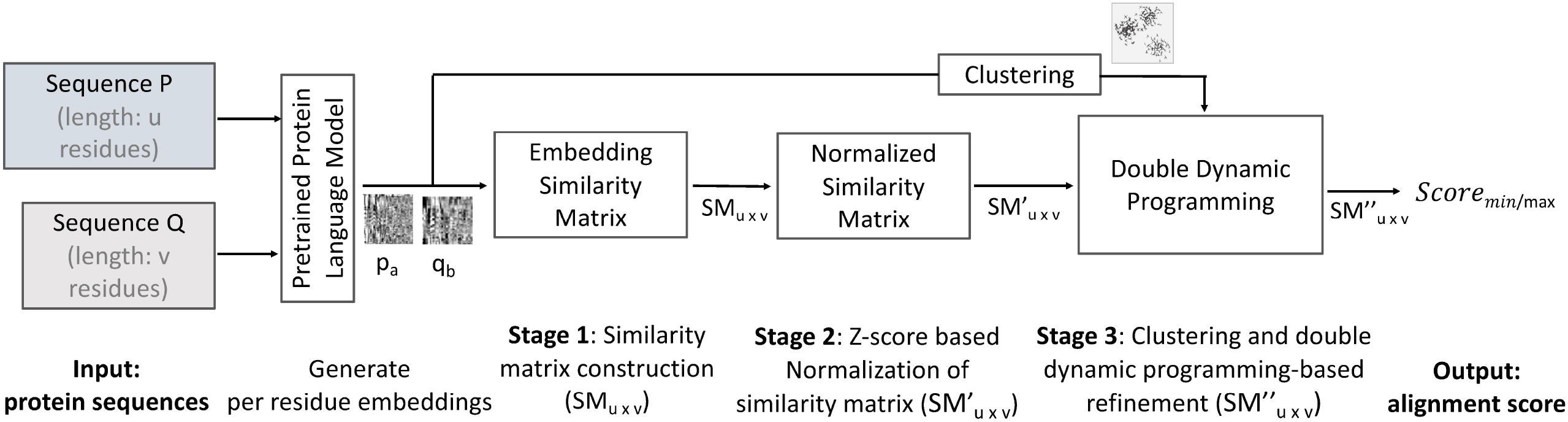
Overview of our approach. Given a pair of input protein sequences, P (of length u) and Q (of length v), residue-level embeddings are generated using a pretrained protein language model. In **Stage 1**, a residue–residue similarity matrix (SM_u×v_) is computed. To reduce noise in the initial similarity matrix, **Stage 2** applies Z-score normalization independently across rows and columns, resulting in a normalized similarity matrix (SM′_u×v_). **Stage 3** further refines the similarity matrix by first performing a Needleman– Wunsch dynamic programming, followed by applying K-means clustering on the combined embeddings from both sequences. Cluster-based weights are then incorporated to adjust similarity scores, and a second Needleman–Wunsch dynamic programming is performed on the final similarity matrix (SM′′_u×v_). Finally, the alignment score derived from this matrix is normalized using either the shorter (Score_min_) or longer (Score_max_) sequence length.

While these embeddings offer a powerful representation of protein sequences, a common practice is to average the residue-level embeddings to obtain a fixed-size vector for each protein^31–33^. This enables fast sequence comparison using distance metrics such as Euclidean distance. While this approach can capture high-level similarities between proteins, it often struggles with accurate residue-level alignment^33,36^.

### Stage 1: Construction of Embedding Similarity Matrix

To address this limitation, we construct a residue–residue similarity matrix that captures the fine-grained spatial relationships between sequences. Specifically, to compare two protein sequences P and Q, with lengths u and v, we first compute an embedding similarity matrix (*SM*_u x v_), where each entry represents the similarity between a pair of residues. The similarity score between residue *a* in sequence P and residue b in sequence Q is computed as:

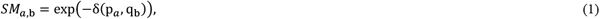

where p_*a*_and q_b_ are the residue-level embeddings of residues *a* (∈P) and b (∈Q), respectively, and δ denotes the Euclidean distance.

### Stage 2: Z-Score-Based Normalization of the Similarity Matrix

To reduce noise, inspired by prior work^36^, we transform the initial similarity matrix (*SM*_u x v_) using a Z-score normalization strategy. For each residue *a* ∈P, we compute the row-wise mean *μ*_*r*_(*a*) and standard deviation *σ*_*r*_(*a*) as follows:

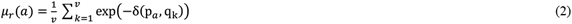

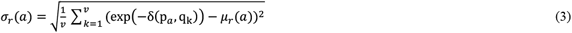

Similarly, for each residue b ∈Q, we compute the column-wise mean *μ*_*c*_(*b*) and standard deviation *σ*_*c*_(*b*) as follows:

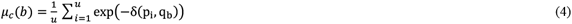

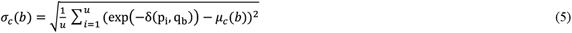

The Z-scores are computed with respect to both the row and column distributions as follows:

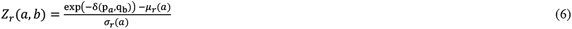

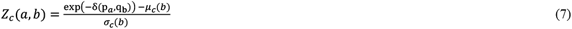

where *Z*_*r*/*c*_(*a, b*) is the row-/column-wise Z-score for a residue pair (*a, b*). The Z-score-based similarity matrix 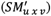 is then obtained by averaging the row-and column-wise Z-scores of each residue pair as follows:

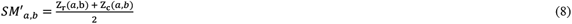

### Stage 3: Refining the Z-score-based similarity matrix with clustering and double dynamic programming

To further remove noise and highlight informative signals, we introduce a new refinement strategy that updates the Z-score–normalized similarity matrix 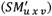 from Stage 2 using clustering and double dynamic programming (DDP). We use Needleman–Wunsch^46^ dynamic programming (with zero gap penalties) on the Z-score-based similarity matrix 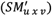. Next, we apply K-means clustering to the combined residue embeddings from both sequences, grouping residues into 20 clusters based on their embedding representations. While this is inspired in part by prior work^47^ that applied clustering in 20-dimensional space for alignment-free protein classification, our use of clustering is fundamentally different, as it refines residue-level similarity matrices for sequence alignment. Based on the resulting cluster assignments, a weight matrix is created to assign higher similarity scores to residue pairs that belong to the same cluster. Specifically, each aligned residue pair is used to adjust the corresponding similarity score by blending it with a cluster-informed value. The updated similarity score between residue *a* (∈P) and residue *b* (∈Q) is computed as:

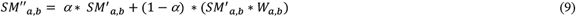

where *α* is a fixed blending factor (*α* = 0.8) and *W*_*a,b*_=1.0 if residues *a* and *b* share the same cluster, and *W*_*a,b*_= 0.5 otherwise. Residue pairs that were not aligned are left unchanged in this step. On this refined similarity matrix 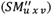, a second Needleman–Wunsch dynamic programming is used to produce the final alignment. Similar refinement strategy is originally proposed in ^48^ and later adopted for contact (or distance)-assisted protein structure prediction^13,14,49^; in contrast, our approach is quite different. Finally, the resulting alignment score (S_align_) is normalized using either the minimum or maximum sequence length to allow fair comparison across protein pairs:

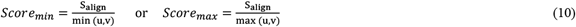

This asymmetric normalization ensures that alignment scores are interpretable across varying sequence lengths, inspired by previous works^20,35,36^.

### Benchmark datasets, methods to compare, and performance evaluation

To evaluate the performance in structural similarity analysis, we benchmark our approach using protein pairs from the PISCES^40^ dataset, containing 19 599 protein pairs with a maximum pairwise sequence similarity of 30% and a minimum chain length of 75 residues. While these protein pairs show detectable homology with an e-value threshold of 10^−4^ using HHsearch^50^, the maximum sequence similarity cutoff of 30% indicates distant homology in the twilight zone. On this dataset, our approach is compared against state-of-the-art protein embedding-based approaches, such as EBA^36^, ProtTucker^31^, TM-Vec^29^, and pLM-BLAST^51^, as well as traditional sequence alignment methods, such as HH-align^50^ and Needleman-Wunsch^46^. Notably, EBA aligns protein sequences in embedding space using dynamic programming on residue-level representations from language models. ProtTucker calculates similarity via Euclidean distance between contrastively trained protein embeddings, while TM-Vec directly predicts TM-scores from embeddings using deep learning. pLM-BLAST identifies local similarity by comparing contextual embeddings. Traditional competing methods include HHalign, which performs profile–profile alignments using hidden Markov models, and Needleman–Wunsch global alignment with a standard BLOSUM^52^ substitution matrix. While our approach also uses Needleman–Wunsch, the purpose of including Needleman-Wunsch with a standard BLOSUM substitution matrix as a competing method is to highlight the performance gap between matrix-based alignments and embedding-enhanced strategies, particularly in low-homology settings. Furthermore, the performance of each method is evaluated by computing the Spearman correlation between its predicted similarity score and the TM-score computed by TM-align^20^, which serves as the ground truth. Correlations were calculated using TM-scores (by TM-align) normalized by the lengths of both the shorter and the longer protein in each pair. It is noted that the reported Spearman correlation values for all competing methods on PISCES dataset are obtained from the reported results of ^36^. To make a fair comparison, we use the same embedding types as competing embedding-based methods.

To further evaluate the performance in transferring CATH domain annotations, we use the same lookup and test set as used by prior work^31,36^. In particular, while the look up set maintains very low sequence similarity to the test set (HVAL < 0), it is ensured that for each protein in the test set, there exists at least one protein in the lookup set with an identical CATH classification at the specified level. On this dataset, our approach is compared against state-of-the-art pLM-based approaches, such as EBA, ProtTucker, and TM-Vec, as well as non pLM-based approaches such as Foldseek^45^, HMMER^2^, and MMseq2^53^. While HMMER and MMseq2 are widely used sequence profile-based approaches, Foldseek is a state-of-the-art structure-based approach. Similar to prior work^31,36^, the performance is evaluated using accuracy as follows:

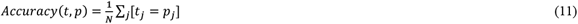

where N denotes the total number of test samples, *t*_*j*_and *p*_*j*_are the ground truth (experimental annotation) and prediction for protein *j*, respectively. [*t*_*j*_= *p*_*j*_] is 1 when the prediction matches the ground truth and 0 otherwise. It is noted that accuracy scores of competing methods are obtained from the reported results of ^31,36^. Notably, both our approach and EBA use ProstT5^31^ embeddings and normalize alignment scores by the length of the longer protein in each aligned pair. In contrast, ProtTucker and TM-Vec rely on embeddings from ProtT5^30^.

## Results and Discussions

### Performance on PISCES dataset

We evaluate the performance of our approach in structural similarity analysis on PISCES dataset, which includes 19,599 protein pairs filtered to ensure low sequence identity (≤30%) and a minimum chain length of 75 residues. **Table 1** presents the Spearman correlation between predicted similarity scores of competing methods and TM-align–computed TM-scores. Spearman correlations are reported with respect to TM-scores normalized by the shorter sequence length (TM_min_) and the longer sequence length (TM_max_), providing a dual perspective on alignment quality. It is worth mentioning that the Spearman correlations of competing methods are obtained from published results reported in ^36^. To make a fair comparison, we use the same embedding types as competing methods.

**Table 1.**
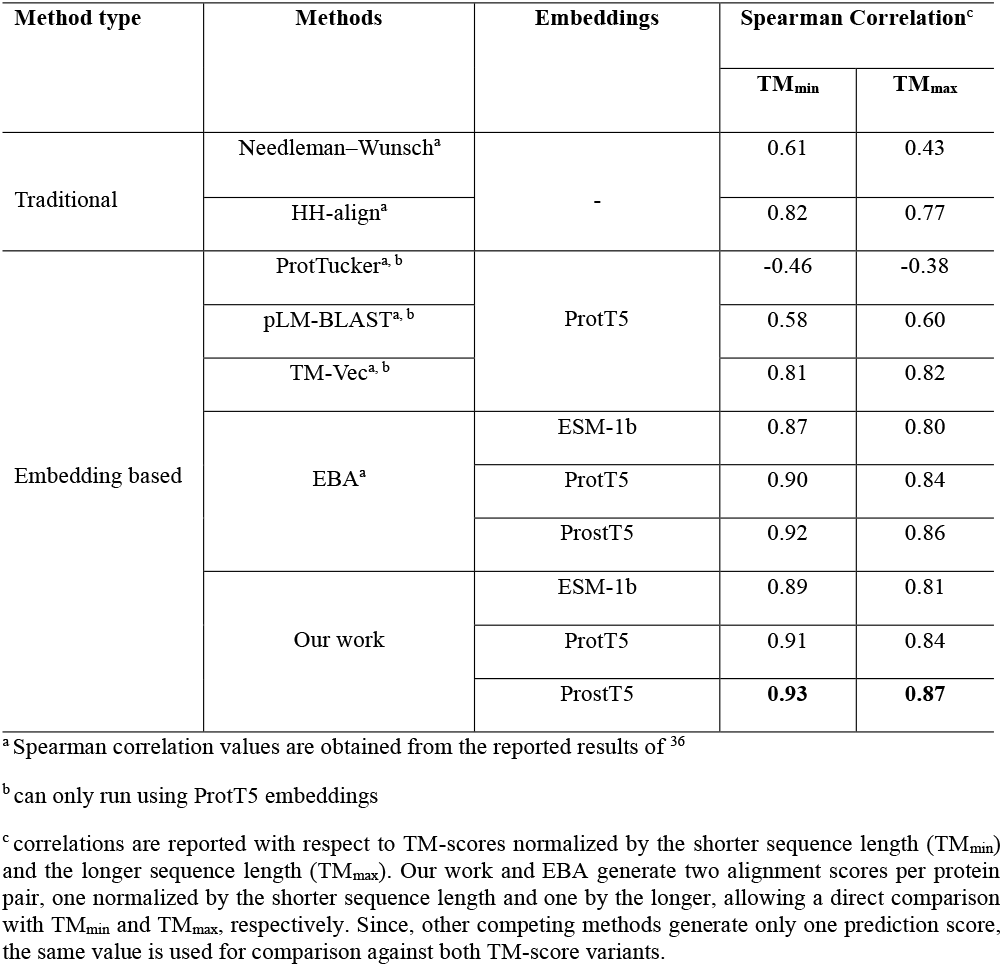
Performance comparison on PSICES dataset (best performance is listed in bold).

As shown in **Table 1**, our approach (refer to Our work), which integrates clustering- and DDP-based refinement of the embedding similarity matrix, consistently achieves the highest Spearman correlations with TM-scores (the ground truth) across all three protein language models: ProstT5, ProtT5, and ESM-1b. Among these, ProstT5 embeddings yield the highest correlation, with Spearman correlations of 0.93 / 0.87 (TM_min_/TM_max_), outperforming all competing embedding-based and traditional methods. Similarly strong correlations are observed with ProtT5 (0.91 / 0.84) and ESM-1b embeddings (0.89 / 0.81), highlighting the effectiveness of our approach across embedding types. Moreover, compared to next best method, EBA, which applies dynamic programming alignment in embedding space, our approach (refer to Our work) achieves higher correlation across all embeddings. For instance, with ProstT5 embeddings, EBA achieves Spearman correlations of 0.92 / 0.86, falling short of the correlations achieved by our approach (0.93 / 0.87). Improvements in Spearman correlations are also noted with ProtT5 (0.91 / 0.84 vs. 0.90 / 0.84) and ESM-1b (0.89 / 0.81 vs. 0.87 / 0.80) embeddings. These improvements highlight the effectiveness of our clustering- and DDP-based approach over the alignment-based state-of-the-art approach, EBA, in remote homology detection. In addition, among other embedding-based methods, TM-Vec, which trains twin neural networks on ProtT5 embeddings to directly predict TM-scores, achieves Spearman correlations of 0.81 / 0.82 (TM_min_/TM_max_) and performs better than some embedding-based and traditional methods, but still lags significantly behind our approach. pLM-BLAST, which uses local alignment heuristics on contextual embeddings, achieves moderate correlations (0.58 / 0.60). ProtTucker, which employs contrastive learning over ProtT5 embeddings, yields negative correlations (–0.46 / –0.38), suggesting its embedding space is not aligned with TM-based structural similarity. Notably, these three approaches—TM-Vec, pLM-BLAST, and ProtTucker—are trained using ProtT5 embeddings and cannot be directly applied across other embedding models, unlike our approach and EBA. Furthermore, while traditional sequence-based tools that do not utilize protein embeddings typically underperform in low-homology, HH-align, which leverages HMM-based profile–profile comparisons, achieves Spearman correlations of 0.82 / 0.77, outperforming several embedding-based methods such as pLM-BLAST and ProtTucker. Although it is outperformed by our approach (refer to Our work) and EBA, HH-align’s performance highlights the enduring value of evolutionary profiles in identifying remote homologs. In contrast, Needleman–Wunsch alignment using a BLOSUM62 matrix performs considerably worse (0.61 / 0.43), illustrating the limitations of fixed substitution scoring. Notably, our approach also uses the Needleman–Wunsch algorithm for global alignment but replaces the BLOSUM matrix with a similarity matrix derived from protein embeddings and refined through clustering and DDP, illustrating the effectiveness of our embedding based scoring matrix. Overall, **Table 1** demonstrates the superior performance of our approach among both traditional and embedding-based approaches on the PISCES dataset, with ProstT5 embeddings yielding the highest Spearman correlations. The consistent improvements across different pLMs and TM-score variants illustrate the effectiveness of our approach in detecting remote homologs.

To further evaluate the predictive quality of our alignment scores, **Figure 2** shows a head-to-head comparison of our predicted scores (refer to Equation 10) using ProstT5 embeddings and the TM-scores computed by TM-align, which serve as the ground truth. Specifically, as shown in **Figure 2**, our predicted alignment scores normalized by the shorter/longer sequence length (Score_min_ / Score_max_) is compared to TM-scores normalized similarly (TM_min_ / TM_max_). In both cases, our approach demonstrates strong positive correlations with the TM-scores, consistent with the high Spearman coefficients reported in **Table 1**. Notably, the gray dashed line in each panel represents a TM-score of 0.5, the commonly used threshold for correctly identifying the correct fold. Overall, this figure demonstrates that our alignment scores, derived from clustering- and DDP-refined embedding similarity matrices, are well-aligned with the structural similarity metric and effective in detecting remote homologs.

**Figure 2.**
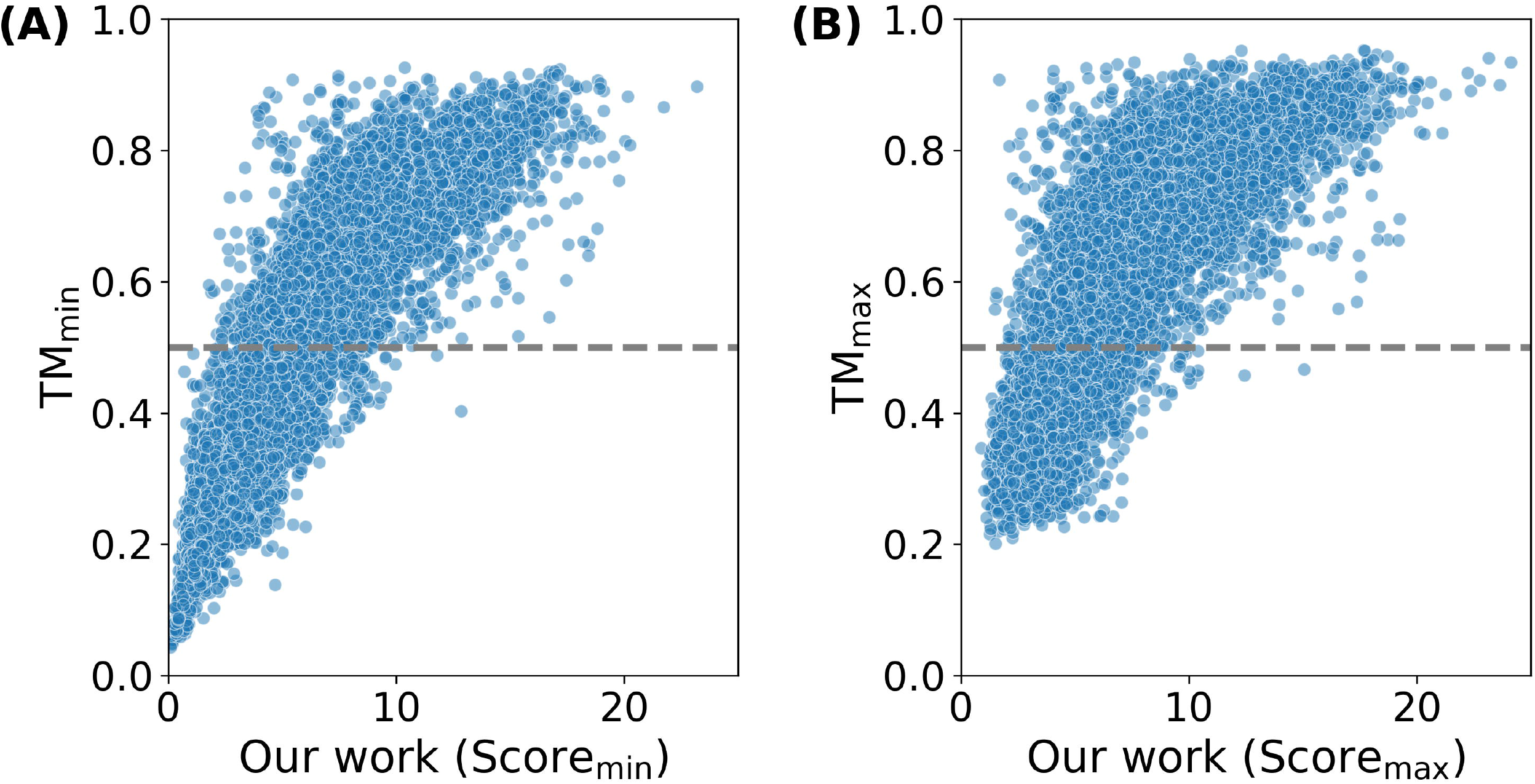
A head-to-head comparison between alignment scores predicted by our approach (using ProstT5 embeddings) and TM-scores (ground truth) computed by TM-align on the PISCES dataset. (**A**) Our predicted alignment scores normalized by the shorter sequence length (Score_min_) vs. TM-scores normalized similarly (TM_min_). (**B**) Our predicted alignment scores normalized by the longer sequence length (Score_max_) vs. TM-scores normalized accordingly (TM_max_). Each data point represents a protein pair from the PISCES dataset, filtered for low sequence identity (≤30%) and a minimum chain length of 75 residues. The horizontal gray dashed line at a TM-score of 0.5 indicates the correct folds.

To complement the structural benchmarking, we also assess the runtime efficiency of our approach (refer to Our work) compared to TM-align. Notably, the reported runtime of our approach excludes the time required to generate residue level embeddings, thereby reflecting only the time required for alignment on a CPU. As shown in **Figure 3**, our approach exhibits a substantial computational advantage over TM-align while maintaining strong structural alignment performance. Specifically, across all tested protein pairs from the PISCES dataset, our approach achieves an average runtime of 0.044 seconds per alignment, compared to 0.172 seconds required by TM-align (refer to **Supplementary Table S1**), representing approximately a fourfold speed-up.

**Figure 3.**
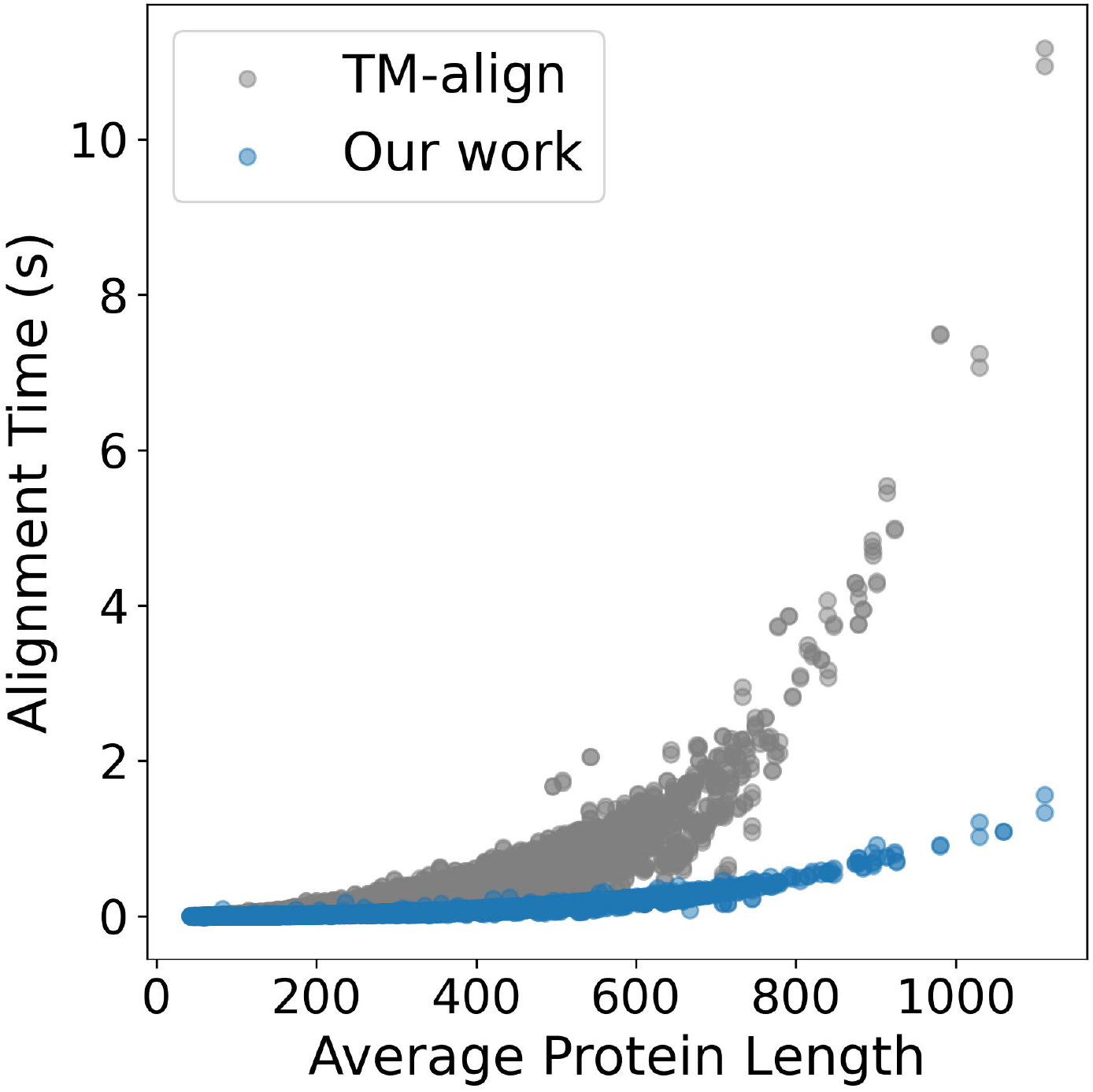
Runtime comparison between our approach and TM-align on the PISCES dataset. X-axis represents average protein length and Y-axis represents alignment runtime (in seconds). Runtime for our approach (blue points) is computed assuming that residue-level embeddings were pre-generated, reflecting only the time required for alignment after embeddings are generated. TM-align’s runtime (gray points) reflects the full computation time of structural alignment. Each data point represents one protein pair from the PISCES dataset.

### Ablation Study

To investigate the individual contributions of each stage in our approach, we perform an ablation study using the PISCES dataset, evaluating Spearman correlations between predicted alignment scores and TM-scores (by TM-align) across three embedding sources: ProstT5, ProtT5, and ESM-1b. As shown in **Table 2**, We evaluate four variants: (1) *Our work*: the full pipeline; (2) *Our work w/o Stage 3*: excludes clustering-and DDP-based refinement; (3) *Our work w/o Stage 2*: excludes Z-score normalization; (4) *Our work w/o Stage 1*: uses averaged residue embeddings to compute Euclidean distance between protein pairs. Notably, similarity scores are expected to exhibit a positive correlation, whereas distance yields a negative correlation.

**Table 2.**
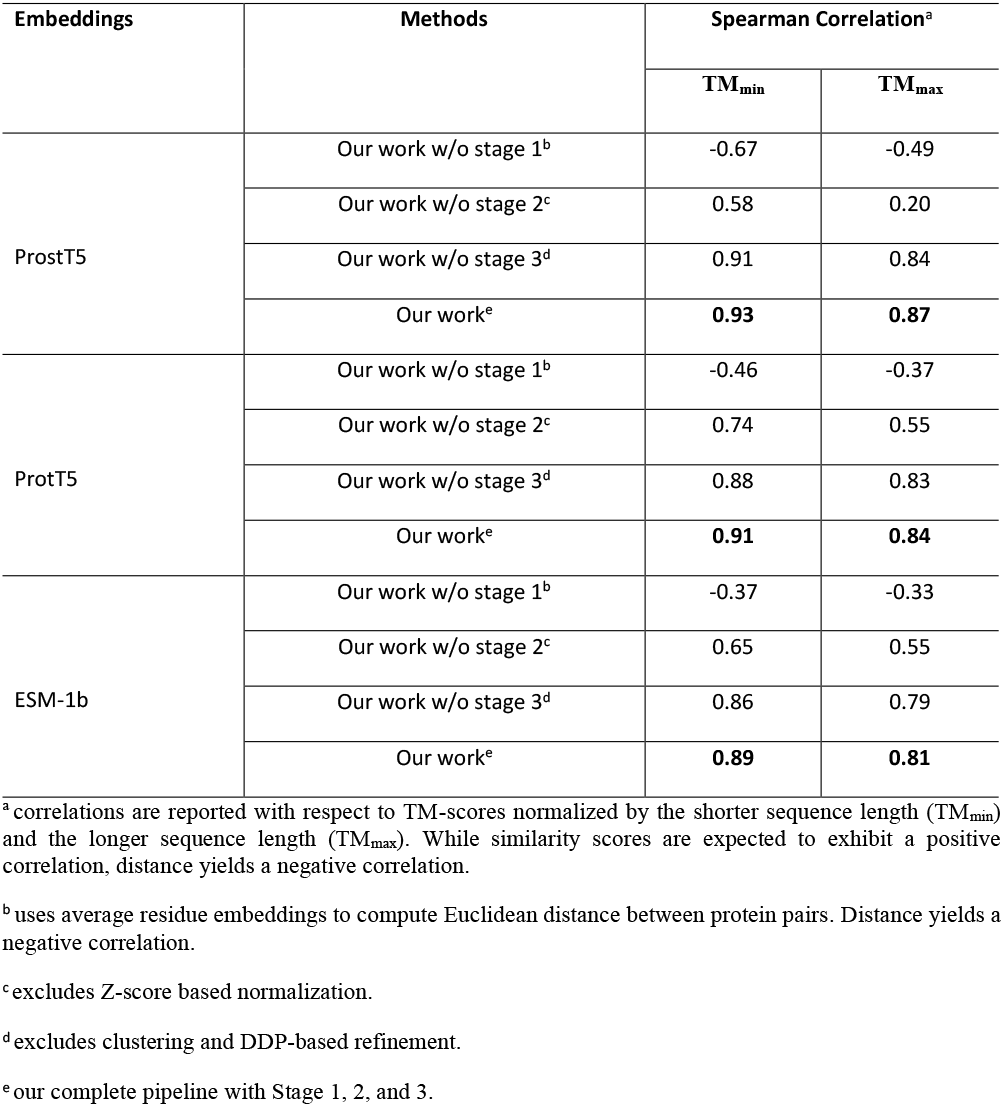
Contribution of individual ablated variants on PISCES dataset (best performance for each embedding is listed in bold).

| Embeddings | Methods | Spearman Correlation<sup>a</sup> |  |
| --- | --- | --- | --- |
|  |  | TM<sub>min</sub> | TM<sub>max</sub> |
| ProstT5 | Our work w/o stage 1<sup>b</sup> | -0.67 | -0.49 |
|  | Our work w/o stage 2<sup>c</sup> | 0.58 | 0.20 |
|  | Our work w/o stage 3<sup>d</sup> | 0.91 | 0.84 |
|  | Our work<sup>e</sup> | **0.93** | **0.87** |
| ProtT5 | Our work w/o stage 1<sup>b</sup> | -0.46 | -0.37 |
|  | Our work w/o stage 2<sup>c</sup> | 0.74 | 0.55 |
|  | Our work w/o stage 3<sup>d</sup> | 0.88 | 0.83 |
|  | Our work<sup>e</sup> | **0.91** | **0.84** |
| ESM-1b | Our work w/o stage 1<sup>b</sup> | -0.37 | -0.33 |
|  | Our work w/o stage 2<sup>c</sup> | 0.65 | 0.55 |
|  | Our work w/o stage 3<sup>d</sup> | 0.86 | 0.79 |
|  | Our work<sup>e</sup> | **0.89** | **0.81** |

<sup>a</sup> correlations are reported with respect to TM-scores normalized by the shorter sequence length (TM<sub>min</sub>) and the longer sequence length (TM<sub>max</sub>). While similarity scores are expected to exhibit a positive correlation, distance yields a negative correlation.
<sup>b</sup> uses average residue embeddings to compute Euclidean distance between protein pairs. Distance yields a negative correlation.
<sup>c</sup> excludes Z-score based normalization.
<sup>d</sup> excludes clustering and DDP-based refinement.
<sup>e</sup> our complete pipeline with Stage 1, 2, and 3.

Across all three types of embeddings, the full pipeline (refer to Our work), which incorporates all three stages, consistently outperforms the other ablated variants by yielding the highest Spearman correlations with TM-scores. Notably, the best performance is observed with ProstT5 embeddings, where the spearman correlations of our full pipeline (refer to Our work) are 0.93/0.87 (TM_min_/TM_max_), followed by ProtT5 (0.91/0.84) and ESM-1b (0.89/0.81) embeddings, illustrating that ProstT5 provides more robust embeddings for remote homology detection. Furthermore, when Stage 3 is removed (refer to Our work w/o Stage 3), which excludes the clustering- and DDP-based refinement introduced in this work, performance decreases consistently across all embeddings. For instance, with ProstT5 embeddings, the correlation drops from 0.93/0.87 to 0.91/0.84. Similar reductions are observed with ProtT5 (from 0.91/0.84 to 0.88/0.83) and ESM-1b embeddings (from 0.89/0.81 to 0.86/0.79). These results indicate that the clustering- and DDP-based refinement introduced in this work provides an additive improvement by refining residue-level similarities beyond what normalization alone can achieve. Moreover, the removal of Stage 2 (refer to Our work w/o Stage 2), which eliminates Z-score normalization, leads to a substantial performance decline across all embeddings. For example, the correlation with ProstT5 embeddings drops sharply to 0.58/0.20, confirming that normalization is essential for denoising the initial similarity matrix. Similar trends are observed with ProtT5 (0.74/0.55) and ESM-1b embeddings (0.65/0.55), demonstrating the importance of denoising the initial similarity matrix. Lastly, using embedding distance alone (refer to Our work w/o Stage 1) yields a negative correlation (–0.67 / –0.49) with ProstT5 embeddings. Similar trends are also observed with ProtT5 (–0.46 / –0.37) and ESM-1b (–0.37/–0.33) embeddings, indicating that raw distances are insufficient for capturing structural similarity and highlights the importance of alignment-based similarity scoring.

Together, these results demonstrate the complementary contributions of each stage in our approach. Specifically, the poor correlations observed when Stage 1 is removed confirm that alignment-based similarity scoring is essential; relying solely on averaged embedding distances fails to capture the structural relationships needed for effective alignment. Stage 2, which applies Z-score normalization, plays a critical role in denoising the initial similarity matrix. Its removal leads to a sharp decline in performance, indicating that normalization is important to amplify informative signals while suppressing background noise. Finally, Stage 3, the clustering- and DDP-based refinement introduced in this work, provides consistent improvements across all embedding types, even after Z-score normalization, demonstrating its effectiveness in refining residue-level similarities. While its contribution is moderate compared to Stage 2, Stage 3 remains a key enhancement that improves alignment quality in an unsupervised manner and strengthens the overall effectiveness of our embedding-based approach for remote homology detection.

Furthermore, we evaluate the average alignment runtime for each ablated version of our approach to assess computational efficiency. It is worth mentioning that all runtime estimates of all ablated variants are computed assuming that residue-level embeddings have been pre-generated, thus reflecting only the time required for alignment on a CPU. As shown in **Supplementary Table S1**, the variant using only embedding distances without alignment (refer to Our work w/o Stage 1) is the fastest, requiring only 7 × 10^−5^ seconds per protein pair, but as discussed earlier, it yields the worst performance. Our work w/o Stage 2 and Our work w/o Stage 3 slightly increase the runtime to 0.020 and 0.021 seconds per pair, respectively, while the full pipeline (refer to Our work), which includes clustering- and DDP-based refinement, requires 0.044 seconds on average. Notably, despite this increase, our full method remains approximately 4 times faster than TM-align. These results suggest that our approach provides an efficient and robust framework for structure-aware sequence alignment.

### CATH Annotation Transfer Performance

To evaluate how well our approach captures functionally relevant structural information from protein sequences, we benchmark its performance on the CATH annotation transfer task. This task involves predicting the CATH classification of a protein at four hierarchical levels—Class (C), Architecture (A), Topology (T), and Homology (H)—by transferring annotations from its most similar protein in a lookup set. Following the protocol used in prior work^31,36^, the lookup and test sets share no detectable sequence similarity (HVAL < 0), ensuring that any successful transfer reflects structural, not sequence, similarity. The accuracy scores of competing methods are obtained from published results reported in ^31,36^.

As shown in **Figure 4**, our approach (refer to Our work) outperforms all competing methods across all four CATH levels and achieves the highest accuracy overall. At the Class level, our approach (refer to Our work) reaches 91% accuracy, similar to the next best method, EBA, but exhibits consistent improvements at other levels over EBA: 85% for Architecture (vs. 84% for EBA), 80% for Topology (vs. 78% for EBA), and 89% for Homology (vs. 88% for EBA). The performance improvements of our approach (refer to Our work) over EBA are consistent across the hierarchy, indicating the effectiveness of our clustering- and DDP-based refinement of similarity matrix in the performance. Notably, both of these approaches use ProstT5 embeddings and normalize alignment scores by the length of the longer protein in each pair, ensuring a fair comparison. While competing pLM-based methods like ProtTucker and TM-Vec use ProtT5 embeddings, our approach (refer to Our work) outperforms these approaches across the CATH hierarchy. While ProtTucker and TM-Vec perform competitively at higher levels, achieving accuracy of 88% and 89% at the Class level and 82% and 83% at the Architecture level, respectively, they exhibit a noticeable decline in performance at finer-grained levels. Specifically, at the Topology level, their accuracy drops to 68% (ProtTucker) and 71% (TM-Vec), falling behind our approach (refer to Our work) by 12 and 9 percentage points, respectively. Similar performance gaps are observed at the Homology level, with ProtTucker and TM-Vec reaching accuracy of 79% and 81%, respectively. Among non–pLM-based methods, Foldseek stands out as the best-performing approach. It achieves accuracy of 77% at Class (Δ14% vs. Our work), 73% at Architecture (Δ12% vs. Our work), 59% at Topology (Δ21% vs. Our work), and 77% at Homology (Δ12% vs. Our work). Similar or more pronounced performance gap can be noted for other non pLM-based methods, HMMER and MMseqs2.

**Figure 4.**
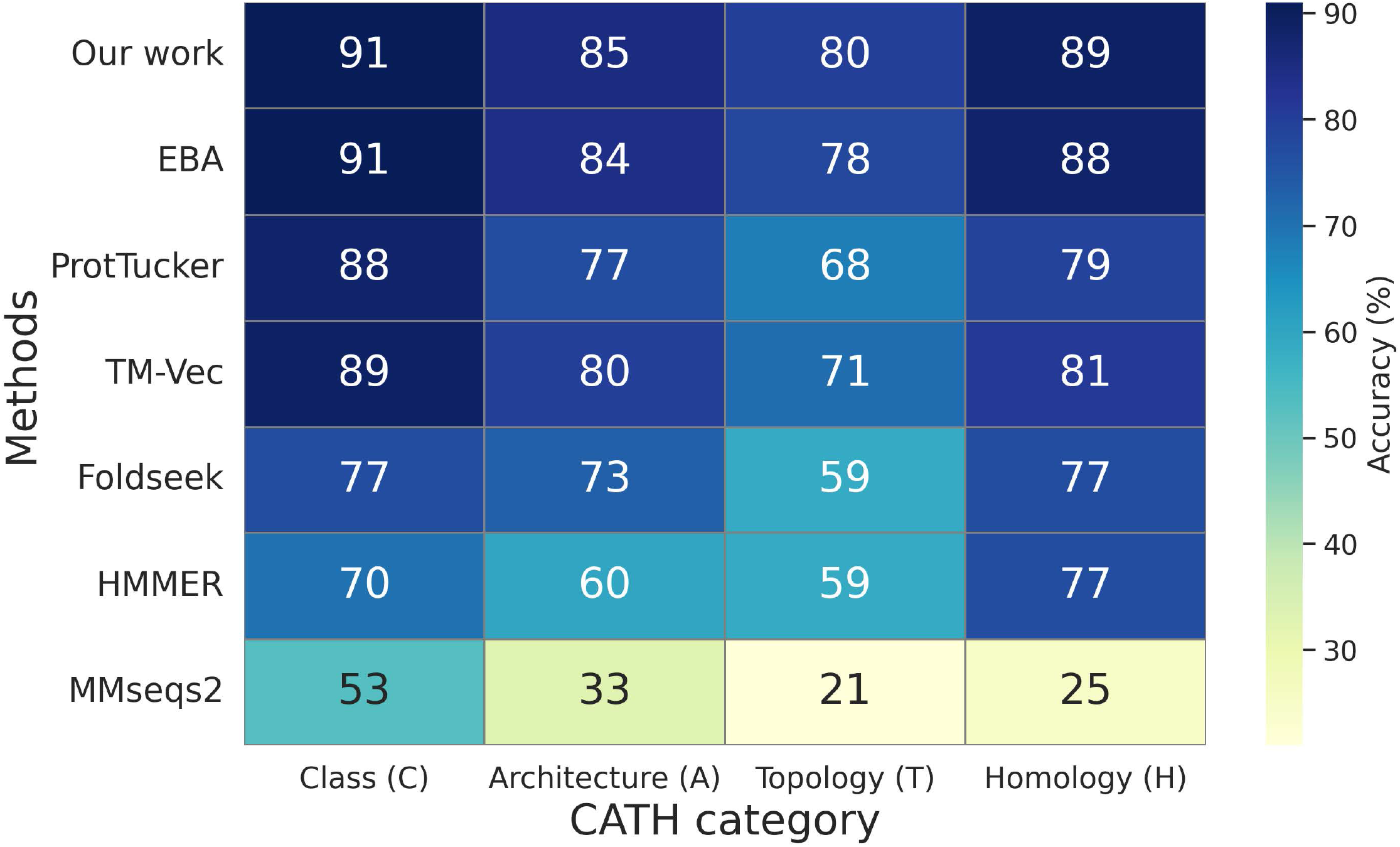
Performance comparison on CATH annotation transfer across four levels: Class (C), Architecture (A), Topology (T), and Homology (H). The heatmap shows accuracy percentages achieved by our clustering- and DDP-refined embedding-based work using ProstT5 embeddings (refer to Our work), compared to other state-of-the-art protein language model (pLM)–based approaches (EBA, ProtTucker, TM-Vec) and non–pLM-based approaches (Foldseek, HMMER, and MMseqs2). Each cell represents the accuracy at the corresponding CATH level, with darker colors indicating higher accuracy. It is noted that accuracy scores of competing methods are obtained from the reported results of ^31,36^. Both our approach and EBA use ProstT5^31^ embeddings and normalize alignment scores by the length of the longer protein in each aligned pair. ProtTucker and TM-Vec rely on embeddings from ProtT5^30^.

Overall, while the performance gap between pLM-based and non pLM-based approaches indicates the effectiveness of pLM embeddings in accurate CATH annotation transfers, the consistent improvements achieved by our approach across all CATH levels highlights the effectiveness of this work over existing state-of-the-art methods, further illustrating the effectiveness of our clustering- and double dynamic programming-based approach in identifying structural and functional relationships across the CATH hierarchy.

## Conclusions

Despite recent advances in protein language models, accurately aligning protein sequences in the twilight zone of sequence similarity remains a major challenge in structural bioinformatics. To address this, we introduce a new unsupervised alignment approach that combines residue-level embeddings from pretrained pLMs with a clustering- and double dynamic programming–based refinement stage. Notably, our approach operates entirely in embedding space and requires no model-specific training, making it broadly applicable across language models and alignment tasks. Moreover, benchmarking on the PISCES dataset demonstrates that our approach achieves the highest Spearman correlation with TM-scores (by TM-align) across all evaluated embedding types, reaching up to 0.93 (TM_min_) and 0.87 (TM_max_) with ProstT5 embeddings, and consistently outperforming both traditional methods such as HH-align and Needleman–Wunsch, and embedding-based state-of-the-art methods including EBA, TM-Vec, and pLM-BLAST, illustrating the effectiveness of the clustering- and DDP-based approach in detecting remote homology. Ablation studies further demonstrate that each component of our pipeline—similarity matrix construction, normalization, and refinement—plays a critical role in driving performance, with clustering- and double dynamic programming-based refinement of embedding similarity matrix introduced in this work show consistent improvements across all embedding types. Moreover, our approach generalizes beyond structural benchmarks: in CATH domain annotation transfer, it achieves highest accuracy across all four classification levels: Class, Architecture, Topology, and Homology—outperforming both pLM-based state-of-the-art methods such as EBA, ProtTucker, and TM-Vec, as well as non–pLM-based methods like Foldseek, HMMER, and MMseqs2. Furthermore, the computational runtime of our approach is still reasonable, making it four times faster than TM-align. Overall, this study demonstrates that our embedding-based alignment refined through unsupervised clustering and double dynamic programming provides a powerful, scalable approach for remote homology detection, with potential to evolve further as protein language models continue to advance.

## Acknowledgement

This work is partially supported by the NSF grant 2435093, and was made possible in part by a grant of high performance computing resources and technical support from the Alabama Supercomputer Authority. The authors would like to thank Dr. Ben Okeke, Dr. Olcay Kursun, Sai Prashanthi Pallati and Priscilla Udomprasert for helpful discussions.

## Data availability

PISCES^40^ dataset is publicly available at http://dunbrack.fccc.edu/pisces/ and https://git.scicore.unibas.ch/schwede/eba_benchmark/-/tree/main/pisces/data?ref_type=heads. CATH^34^ dataset is publicly available at https://www.cathdb.info and https://git.scicore.unibas.ch/schwede/eba_benchmark/-/tree/main/cath/data?ref_type=heads.

## Authors’ contributions

R.S.: Writing - review & editing, Writing -- original draft, Software, Visualization, Validation; N.R.: Writing - review & editing, Writing -- original draft, Software, Validation; C.S.: Writing - review & editing, Writing -- original draft, Validation; S.B.: Writing - review & editing, Writing -- original draft, Visualization, Validation, Supervision, Software, Resources, Project administration, Methodology, Investigation, Funding acquisition, Formal analysis, Data curation, Conceptualization. All authors approve the final version.

