## Supplementary Table 1 for "Evaluating the Significance of Embedding-Based Protein Sequence Alignment with Clustering and Double Dynamic Programming for Remote Homology"

**Robert Spicer^1^, Nilanjana Raychawdhary^2^, Cheryl Seals^2^, Olcay Kursun^1^, Ben Okeke^3^, Sutanu Bhattacharya^1,*^**

^1^Department of Computer Science & Computer Information Systems, Auburn University at Montgomery, Montgomery, AL 36117, USA.

^2^Department of Computer Science & Software Engineering, Auburn University, Auburn, AL 36849, USA.

^3^Department of Biology and Environmental Sciences, Auburn University at Montgomery, Montgomery, AL 36117, USA.

*Corresponding author:	Sutanu Bhattacharya

309L Goodwyn Hall, Auburn University at Montgomery,

7400 East Dr, Montgomery, AL 36117, USA

**Table S1: Average computation time on PISCES dataset.** We compute average alignment time for ablated variants of our work and TM-align. In particular, we evaluate four variants: (1) *Our work*: the full pipeline; (2) *Our work w/o Stage 3*: excludes clustering- and double dynamic programming (DDP)-based refinement; (3) *Our work w/o Stage 2*: removes Z-score normalization; (4) *Our work w/o Stage 1*: uses only embedding distance, with no alignment or matrix construction. Notably, the reported runtime of our work excludes the time required to generate residue level embeddings, thereby reflecting only the time required for alignment on a CPU.

|  | Our work w/o stage 1 | Our work w/o stage 2 | Our work w/o stage 3 | Our work | TM-align |
| --- | --- | --- | --- | --- | --- |
| Average alignment time (s) | 7 * 10^-5^ | 0.020 | 0.021 | 0.044 | 0.172 |
